# Mis-regulation of Zn and Mn homeostasis is a key phenotype of Cu stress in *Streptococcus pyogenes*

**DOI:** 10.1101/2022.09.28.509676

**Authors:** YoungJin Hong, Eilidh S Mackenzie, Samantha J Firth, Jack RF Bolton, Louisa J Stewart, Kevin J Waldron, Karrera Y Djoko

**Author notes:** These authors contributed equally to the work (YoungJin Hong, Eilidh S Mackenzie, Samantha J Firth, Jack RF Bolton, Louisa J Stewart). Author order was decided by an on-line list randomiser.

## Abstract

All bacteria possess homeostastic mechanisms that control the availability of micronutrient metals within the cell. Cross-talks between different metal homeostasis pathways within the same bacterial organism have been reported widely. In addition, there have been previous suggestions that some metal uptake transporters can promote adventitious uptake of the wrong metal. This work describes the cross-talk between Cu and the Zn and Mn homeostasis pathways in Group A Streptococcus (GAS). Using a Δ*copA* mutant strain that lacks the primary Cu efflux pump and thus traps excess Cu in the cytoplasm, we show that growth in the presence of supplemental Cu promotes downregulation of genes that contribute to Zn or Mn uptake. This effect is not associated with changes in cellular Zn or Mn levels. Co-supplementation of the culture medium with Zn or, to a lesser extent, Mn alleviates key Cu stress phenotypes, namely bacterial growth and secretion of the fermentation end-product lactate. However, neither co-supplemental Zn nor Mn influences cellular Cu levels or Cu availability in Cu-stressed cells. In addition, we provide evidence that the Zn or Mn uptake transporters in GAS do not promote Cu uptake. Together, the results from this study strengthen and extend our previous proposal that mis-regulation of Zn and Mn homeostasis is a key phenotype of Cu stress in GAS.

**GRAPHICAL ABSTRACT:** 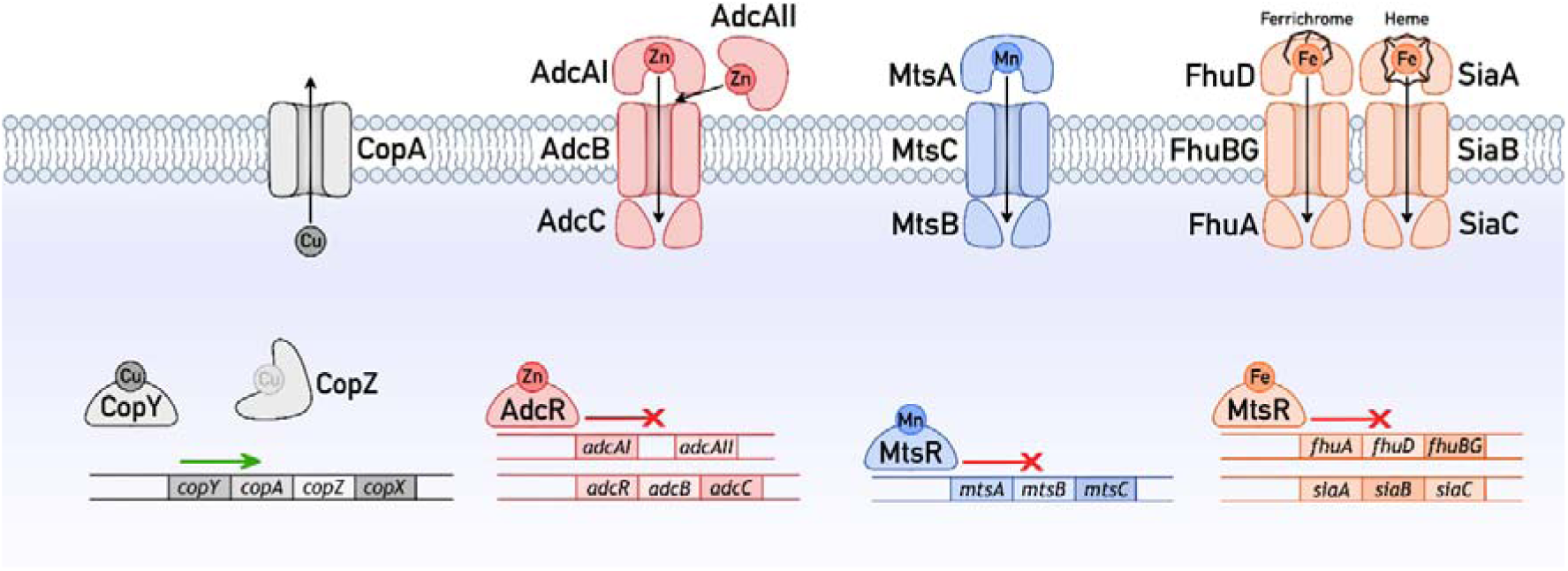

## INTRODUCTION

In general, metal homeostasis systems are specific for their cognate metals. Each metal sensor, importer, exporter, storage protein, and metallochaperone is specialised to manage the cellular availability of their cognate metal ion, and is typically inefficient in managing non-cognate metal ions. However, cross-talks between different metal homeostasis systems can occur. Perturbations to a metal homeostasis system, whether as a result of exposure to an excess of the cognate metal ion, depletion of that metal ion, or genetic manipulation of a component of that system, can lead to the accumulation or depletion of a *different* metal ion in the cell, and/or the transcriptional activation or repression of *another* metal homeostasis system. In prokaryotes, cross-talks between Cu and Fe homeostasis systems^1–5^, Cu and Zn^4,6,7^, Mn and Zn^8,9^, Fe and Zn^10^, and Mn and Fe^11–14^ have been described.

The molecular mechanisms behind such cross-talks and their corresponding cellular outcomes vary. Some metals play *direct* roles in the homeostasis of a different metal. For example, the CopY Cu sensor from *Streptococcus pneumoniae* is Zn-dependent. CopY derepresses expression of Cu efflux genes in response to increases in cellular Cu availability^15^. Conversely, CopY represses expression in response to decreases in Cu availability. However, Zn is required to stabilise the repressor form of this metallosensor^16^. Thus, Zn supplementation suppresses expression of the CopY regulon^15^ while Zn limitation upregulates it, even without additional exposure to Cu^7^.

An excess of a metal ion can bind adventitiously to non-cognate metal homeostasis proteins and interfere *directly* with the function of these proteins. In *Streptococcus pneumoniae*, an excess of Zn can bind adventitiously to the Mn-binding site of the Mn uptake protein PsaA, preventing uptake of Mn *via* the PsaABC Mn-importing ABC transporter, limiting cellular Mn, and subsequently inducing expression of Mn uptake genes^8,9^. An excess of Zn is also thought to bind adventitiously to the Mn- sensing site of the Mn sensor PsaR and promote inadvertent derepression of Mn uptake genes^17^.

Adventitious binding of a metal ion to non-cognate sites can also *indirectly* influence cellular handling of another metal. *Bacillus subtilis* responds to excess Cu by increasing expression of Fe uptake genes^1,5^. In this organism, excess Cu mismetalates Fe-S clusters and thus inactivates Fe-S cluster-dependent enzymes as well as Fe-S cluster assembly machineries^5^. The displaced Fe atoms should have increased cellular Fe availability and thus suppressed (rather than induced) expression of Fe uptake genes *via* the Fe sensor Fur. However, the loss of functional Fe-S clusters transcriptionally induces expression of more cluster assembly machineries^5^. This generates a cellular Fe sink, lowers cellular Fe availability, and thus induces (rather than suppresses) expression of Fe uptake genes.

We previously identified a potential cross-talk between Cu stress and Zn, Mn, and Fe homeostasis in the Gram-positive bacterium *S. pyogenes* (Group A Streptococcus, GAS)^18^. Like other streptococci, GAS employs a single system for Cu sensing and efflux, controlled by the CopY Cu sensor^19^ (Figure 1). This organism is not known to import, use, or store nutrient Cu. When cellular Cu availability rises, CopY transcriptionally derepresses expression of the Cu-effluxing P_1B_-type ATPase CopA, the Cu-binding metallochaperone CopZ, and a putative membrane-associated protein of unknown function named CopX^18,19^. Zn sensing and homeostasis in GAS are composed of two systems, one each for Zn uptake and Zn efflux, which are controlled by the AdcR and GczA Zn sensors, respectively^20,21^. Under conditions of low cellular Zn availability, AdcR transcriptionally derepresses expression of the Zn-importing AdcAI/AdcAII-AdcBC ABC transporter (Figure 1), along with accessory proteins such as the poly-His triad protein Pht. Under conditions of high cellular Zn availability, GczA transcriptionally activates expression of the Zn-effluxing cation diffusion facilitator CzcD. The uptake of Mn and Fe in GAS is controlled by the dual Mn/Fe sensor MtsR^22,23^. In response to low cellular Mn availability, MtsR transcriptionally derepresses expression of the Mn-importing MtsABC ABC transporter (Figure 1). In response to low cellular Fe availability, this sensor derepresses expression of a variety of Fe uptake systems, including the ferrichrome-importing FhuADBG and heme-importing SiaABC ABC transporters (Figure 1). Finally, GAS also employs the cation diffusion facilitator MntE to efflux Mn^24^ and the P_1B_-type ATPase PmtA to efflux Fe^25^, although neither transporter is known to be directly regulated by a Mn- or Fe-sensing transcriptional regulator.

**Figure 1.**
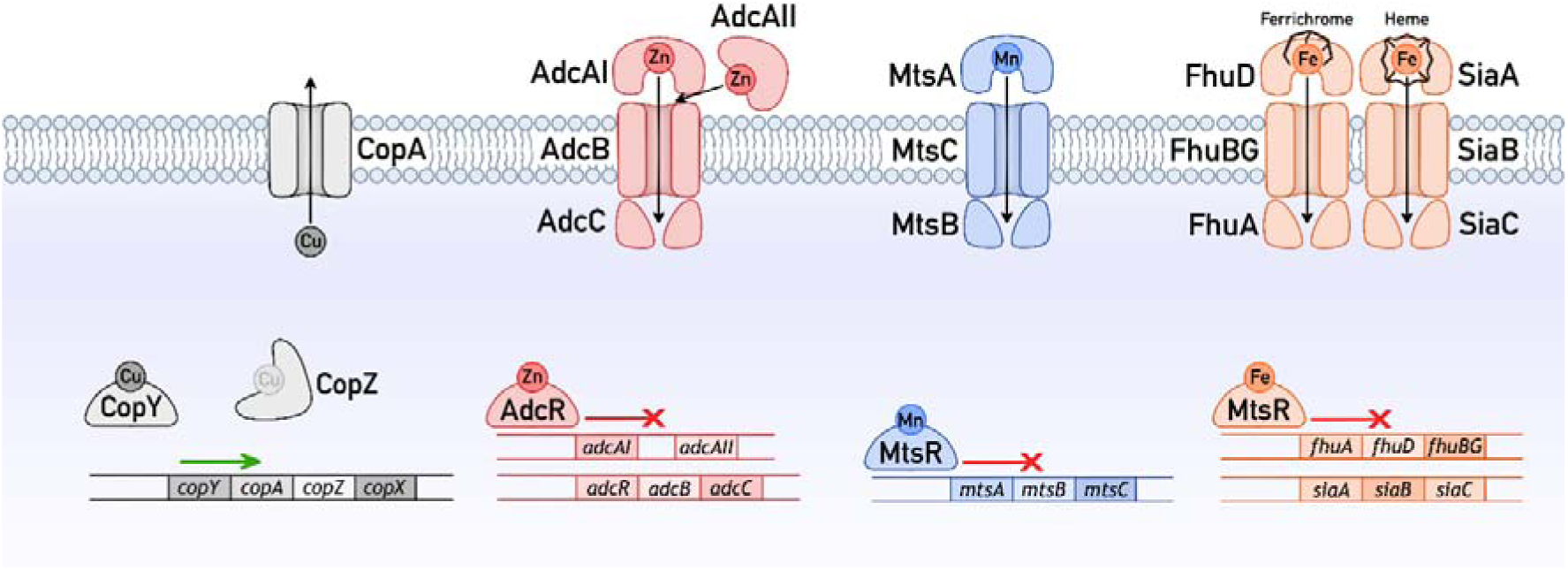
Cu, Zn, Mn, and Fe homeostasis systems in GAS. Only components that are directly relevant to this work are shown. The metallosensor responsible for regulating the transcriptional responses to each metal is shown, along with the direction of transcriptional regulation. Transporters responsible for the efflux of Cu or uptake of Zn, Mn, and Fe are also shown.

Our previous work found that Cu stress in GAS was associated with mis-repression of AdcR- regulated genes, namely *adcAI* and *adcAII*, as well as MtsR-regulated genes, namely those in the *fhu* and *sia* operons^18^. Interestingly, transcription of the *mts* operon, which is also controlled by MtsR, remained unperturbed. Similarly, there was no effect on the expression of *czcD*, *mntE*, or *pmtA*, which are not controlled by AdcR or MtsR. Therefore, the goal of this study is to describe the cross-talks between Cu stress and the AdcR and MtsR regulons in GAS in more detail.

## METHODS

### Data presentation

Except for growth curves, individual data points from independent experiments are plotted, with lines or shaded columns representing the means, and error bars representing standard deviations. Growth curves show the means of independent experiments, with shaded regions representing standard deviations. The number of independent experiments (*N*) is stated in each figure legend. All quantitative data were plotted in GraphPad Prism. Unless otherwise stated, *P* values were calculated by two-way ANOVA using Prism’s statistical package.

### Reagents

All reagents were of analytical grade and obtained from Merck or Melford Chemicals unless otherwise indicated. The sulfate, nitrate, and chloride salts of metal ions were used interchangeably because numerous experiments in the laboratory did not find any meaningful differences between them. All reagents were prepared in deionised water.

### Strains and culture conditions

GAS M1T1 5448 strains (Supporting Table 1) were propagated from frozen glycerol stocks onto solid Todd-Hewitt medium (Oxoid) containing 0.2 w/v % yeast extract (THY) medium without any antibiotics. Unless otherwise indicated, liquid cultures were prepared in a chemically defined medium containing glucose as the carbon source (CDM-glucose^18^). This medium routinely contains <200 nM of total Zn, Cu, or Fe, and <20 nM of total Mn. Catalase (50 µg/mL) was added to all solid and liquid media.

### Bacterial growth

Growth was assessed at 37 °C in flat-bottomed 96-well plates using an automated microplate shaker and reader. Each well contained 200 µL of culture. Each plate was sealed with a gas permeable, optically clear membrane (Diversified Biotech). OD_600_ values were measured every 20 min. The plates were shaken at 200 rpm for 1 min in the double orbital mode immediately before each reading. OD_600_ values were not corrected for path length (*ca*. 0.58 cm for a 200 µL culture).

### ICP MS analyses

GAS was cultured in 40 mL of CDM-Glucose. At the desired time points, cultures were harvested (5,000 x *g*, 4°C, 10 min), washed once with Tris-HCl buffer (50 mM, pH 8.0) containing *D*-Sorbitol (1 M), MgCl_2_ (10 mM), and EDTA (1 mM), and twice with ice-cold PBS. The pellet was resuspended in ice-cold PBS (1 mL). An aliquot was collected for the measurement of total protein content. The remaining suspension was re-centrifuged. The final pellet was dissolved in conc. nitric acid (65 v/v %, 150 µL, 80 °C, 1 h) and diluted to 3.5 mL with deionised water. Total metal levels in these samples were determined by inductively-coupled plasma mass spectrometry (ICP MS) using ^45^Sc, ^69^Ga, and ^209^Bi as internal standards (1 ppb each).

It is important to note that any intact but unviable bacterial cells as well as viable but unculturable cells were harvested together with viable and culturable cells. Therefore, all types of cells contributed to total metal levels as measured by ICP MS. For this reason, total metal levels were normalised to total biomass as measured by cellular protein content (and not to total viable colony forming units).

### Secreted lactate levels

GAS was cultured in 96-well plates as described earlier for growth analysis. After 24 h of growth, samples were centrifuged (5,000 x *g*, 4°C, 10 min) and concentrations of lactate in the supernatants were determined using K-LATE kit (Megazyme).

### GapA activity

Bacteria were cultured in 40 mL of CDM-glucose. After 8 h of growth, bacteria were harvested (5,000 x *g*, 4°C, 10 min), washed once with Tris-HCl buffer (50 mM, pH 8.0) containing *D*-Sorbitol (1 M), MgCl_2_ (10 mM), and EDTA (1 mM), and twice with ice-cold PBS. Bacterial pellets were resuspended in a buffer containing sodium phosphate (100 mM) and triethanolamine (80 mM) at pH 7.4, transferred to a tube containing Lysing Matrix B (MP Biomedicals), and lysed in a FastPrep 24G instrument (MP Biomedicals, 10 m/s, 20 s, 2 cycles). Intact cells and cell debris were removed by centrifugation (20,000 x *g*, 1 min) and cell-free lysate supernatants were kept on ice and used immediately.

To determine GapA activity, the reaction mixture contained NAD^+^ (4 mM), *DL-*glyceraldehyde-3-phosphate (G3P, 0.3 mg/mL), sodium phosphate (100 mM), DTT (1 mM), and triethanolamine (80 mM) at pH 7.4. Each reaction (100 µL) was initiated by addition of cell-free lysate supernatants (10 µL). Absorbance values at 340 nm were monitored for up to 10 min at 37 °C. Initial rates of reaction were normalised to protein content in the cell-free lysate supernatants. Control reactions without any G3P were always performed in parallel. One unit of activity was defined as 1000 nmol NAD^+^ oxidised min^-1^ mg protein^-1^.

### SodA activity

SodA activity was assessed qualitatively using a gel-based assay. First, bacteria were cultured and pelleted as described above for measurements of GapA activity. Cell-free lysate supernatants were also prepared as above, but using Tris-HCl (50 mM, pH 8.0) containing NaCl (150 mM). Protein content in the cell-free lysate supernatants was determined and 8 µg of proteins were resolved on 15% native polyacrylamide gels. The gels were incubated in buffer containing potassium phosphate (50 mM, pH 7.8), EDTA (1 mM), nitro blue tetrazolium chloride (0.25 mM), and riboflavin (0.05 mM), then exposed to light to detect SodA activity. Purified recombinant Fe-loaded and Mn-loaded SodA from *S. pyogenes* (metal-verified by ICP MS; 0.25 µg each) were loaded in parallel as controls. Incubation of replica gels with InstantBlue® Coomassie Protein Stain (Abcam) was performed to assess sample loading. All gels were imaged using a ChemiDoc™ imaging system (Bio-Rad), using the same settings for all gels compared within a single experiment.

### Protein content

Total protein content in all cell extracts was determined using the QuantiPro BCA Assay Kit (Sigma).

### RNA extraction

Bacteria were cultured in 1.6 mL of CDM-glucose. At the desired time points, cultures were centrifuged (4,000 x *g*, 4°C, 5 min). Bacterial pellets were resuspended immediately in 500 µL of RNAPro Solution (MP Biomedicals) and stored at -80°C until further use. Bacteria were lysed in Lysing Matrix B and total RNA was extracted following the manufacturer’s protocol (MP Biomedicals). RNA extracts were treated with RNase-Free DNase I enzyme (New England Biolabs). Complete removal of gDNA was confirmed by PCR using gapA-check-F/R primers (Supporting Table 2). gDNA-free RNA was purified using Monarch RNA Cleanup Kit (New England Biolabs) and visualised on an agarose gel.

### qRT-PCR analyses

cDNA was generated from 1 µg of RNA using the SuperScript® IV First-Strand Synthesis System (Invitrogen). qPCR was performed in 20 µL reactions using Luna qPCR Universal qPCR Master Mix (New England Biolabs), 5 ng of cDNA as template, and 0.4 µM of the appropriate primer pairs (Supporting Table 2). Each sample was analysed in technical duplicates. Amplicons were detected in a CFXConnect Real-Time PCR Instrument (Bio-Rad Laboratories). *C*_q_ values were calculated using LinRegPCR after correcting for amplicon efficiency. *C*_q_ values of technical duplicates were typically within ± 0.25 of each other. *holB*, which encodes DNA polymerase III, was used as the reference gene.

## RESULTS

### Cu stress is associated with mis-regulation of AdcR-and MtsR-controlled genes

We previously showed that growth in a metal-limited, chemically defined medium in the presence of supplemental Cu led to aberrant regulation of metal homeostasis in the GAS 5448 Δ*copA* mutant strain^18^. Specifically, expression of genes under the control of AdcR and MtsR became downregulated, as determined by RNA-seq of the entire transcriptome and qRT-PCR of select genes. These effects appeared after >4 h of growth and correlated with depletion in intracellular glutathione. Our model was that decreasing glutathione levels during bacterial growth led to decreased intracellular Cu buffering capacity, increased Cu availability, and the appearance of multiple Cu stress phenotypes^18^, such as impaired bacterial growth, loss of bacterial viability, decreased production of lactate from fermentation, and the aforementioned mis-regulation of AdcR and MtsR-controlled genes.

In this work, expression of AdcR-and MtsR-regulated genes in Δ*copA* cells was examined beyond 4 h of growth up to 8 h using qRT-PCR. As the control, expression of the Cu-inducible, CopY-regulated gene *copZ* was assessed in parallel. Figure 2A confirms that *copZ* was upregulated at all time points, consistent with the expected increase in intracellular Cu levels and availability in these Cu-treated cells.

**Figure 2.**
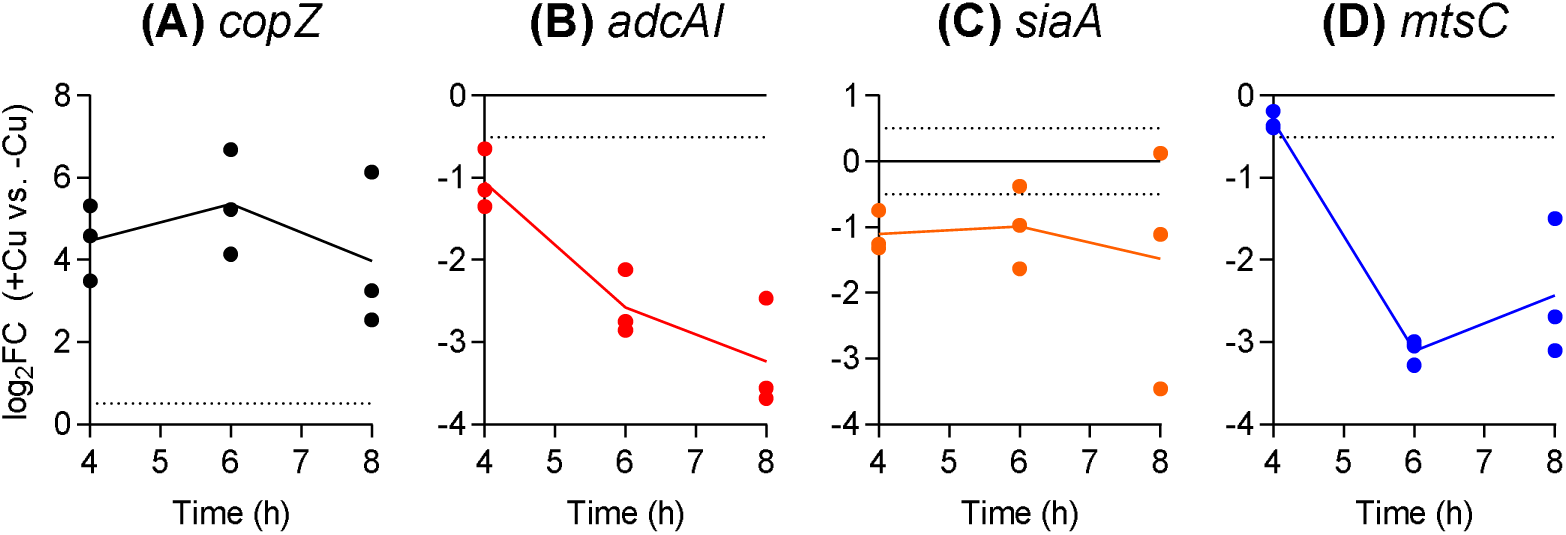
Effects of Cu treatment on expression levels of (A) *copZ*, (B) *adcAI*, (C) *siaA*, and (D) *mtsC.* The GAS 5448Δ*copA* mutant strain was cultured with or without added Cu (5 µM) for *t =* 4, 6, or 8 h (*N* = 4). mRNA levels of each target gene in Cu-supplemented cultures (+Cu) were determined by qRT-PCR and normalised to those in the corresponding unsupplemented samples (-Cu) that were cultured for the same time periods. Dotted horizontal lines represent the sensitivity limit of the assay (log_2_FC = ± 0.5). Data from individual replicates are shown. Columns indicate means. Expression levels of *adcAI* (*P* = 0.021) and *mtsC* (*P* = 0.050) were time-dependent but not those of *siaA* (*P* = 0.87) or *copZ* (*P* = 0.29).

As reported previously, *adcAI* and *adcAII* were downregulated in Cu-treated Δ*copA* cells that were sampled at 4 h (Figure 2B, Supporting Figure 1A). Both genes became further repressed at 6 h and 8 h. Transcription of another AdcR-regulated gene, namely *adcC*, remained relatively unperturbed (Supporting Figure 1B). These results are consistent with differential regulation of the *adc* genes by AdcR^20^ and with our previous report, which detected no change in *adcC* or *adcB* expression in response to Cu treatment^18^. Cu treatment led to downregulation of *siaA* and *fhuA* in Δ*copA* cells that were sampled at 4 h (Figure 2C and Supporting Figure 1C). The effect of Cu treatment on these genes became less clear in cells sampled at 6 and 8 h. Consistent with our previous study^18^, expression of a different MtsR-regulated gene, namely *mtsC*, was not perturbed at 4 h (Figure 2D). However, *mtsC* did become downregulated at 6 and 8 h. Overall, these observations support our previous conclusion that Cu stress is associated with mis-regulation of AdcR and MtsR-controlled genes. However, it is important to note that the effect varies with different genes, consistent with the established action of both metallosensors in differentially regulating their targets^20,26^.

### Cu stress is not associated with changes in cellular Zn, Mn, or Fe

Given their known roles in Zn, Mn, or Fe homeostasis, changes in AdcR-and MtsR-regulated genes may be associated with changes in cellular Zn, Mn, or Fe levels. These genes may become repressed (*i.e.* the effect) in response to increases in cellular Zn, Mn, or Fe levels and/or availability (*i.e.* the cause). Conversely, since the protein products of *adcAI*, *adcAII*, *mtsC*, *fhuA*, or *siaA* contribute to Zn, Mn, or Fe (ferrichrome or heme) uptake^27–30^, repression of these genes (*i.e.* the cause) may lower cellular Zn, Mn, or Fe levels and/or availability (*i.e.* the effect).

In agreement with our previous work^18^, growth in the presence of supplemental Cu increased total cellular Cu levels but did not affect total cellular Zn, Mn, or Fe levels in Δ*copA* cells that were sampled after 4 h of growth (Supporting Figure 2). In an earlier version of this study, we observed a marked decrease in cellular Zn in Δ*copA* cells that were sampled after 8 h of growth^31^. However, reanalysis of the data revealed that Zn levels in the control cells (not treated with any metal) were abnormally high when compared with numerous other Δ*copA* cells from our laboratory that were prepared under identical conditions but measured in separate ICP MS or ICP OES runs. Therefore, we re-measured cellular metal levels in our original Δ*copA* samples, along with two additional independent replicates. As reported in the earlier version of our work, there was an increase in cellular Cu levels after 8 h of growth in the presence of supplemental Cu (Figure 3A), but there was no change in cellular Mn or Fe levels (Figures 3B-C). We noted a large variability in the Fe data, not unlike the variability in the expression patterns of *siaA* and *fhuA* (*cf.* Figure 2C and Supporting Figure 1C). Potential sources for this variability have not been identified. More crucially, contrary to our previous claim, our new results show that Cu treatment for 8 h did not perturb cellular Zn levels in Δ*copA* cells (Figure 3D).

**Figure 3.**
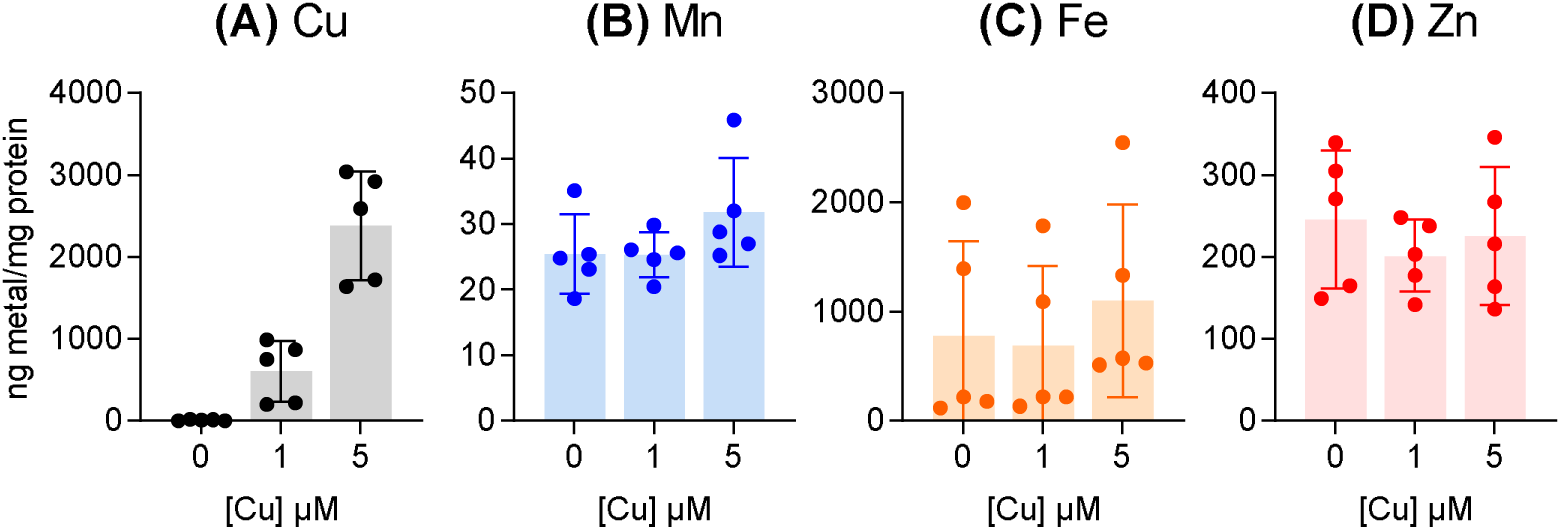
Effects of Cu treatment on cellular levels of (A) Cu, (B) Mn, (C) Fe, and (D) Zn. The GAS 5448Δ*copA* mutant strain was cultured with supplemental Cu (0, 1, or 5 µM) for *t* = 8 h (*N* = 5). Total cellular levels of all metals were measured by ICP MS and normalised to total protein content. Data from individual replicates are shown. Columns indicate means.Error bars represent SD. Cu treatment led to an increase in total cellular Cu (*P* < 0.0001) but not Mn (*P* = 0.33), Fe (*P* = 0.20), or Zn (*P* = 0.68).

As an additional assessment of cellular Mn levels, which were often near the detection limit of our assay, we measured the activity of the superoxide dismutase SodA in Δ*copA* cell-free extracts. SodA from *S. pyogenes* is active with either Mn or Fe in the catalytic site, although enzyme activity with Mn is much higher^32^. Studies with Mn-deficient Δ*mtsABC* mutant strains of GAS indicate that loss of cellular Mn is associated with decreased SodA activity^28,32^. However, growth of the Δ*copA* mutant strain in the presence of Cu for 8 h did not reduce SodA activity in these cells (Figure 4). This result supports our conclusion that Cu stress does not perturb cellular Mn levels.

**Figure 4.**
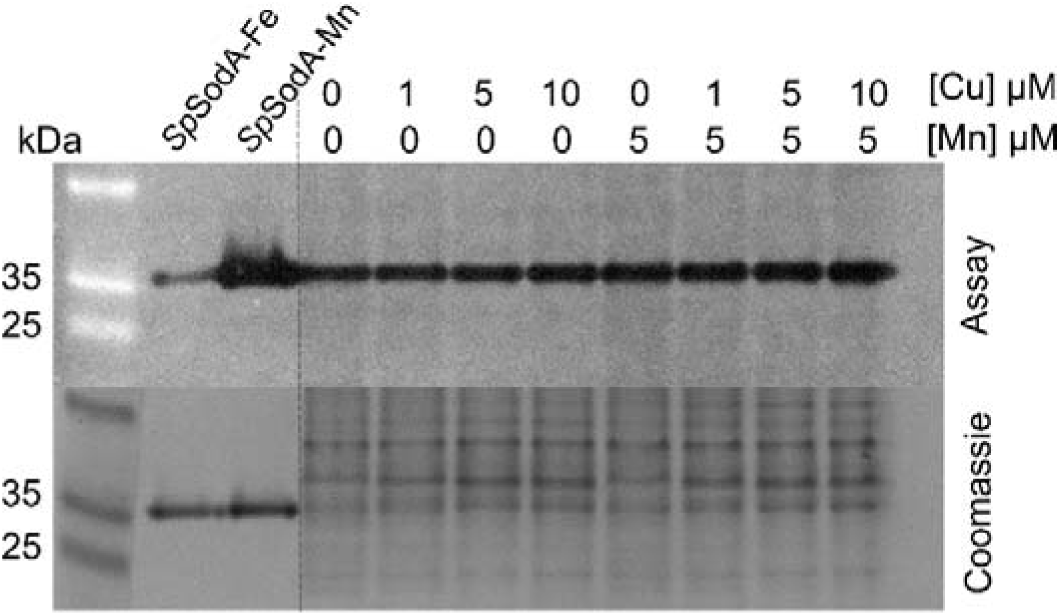
Effects of Cu treatment on SodA activity. The GAS 5448Δ*copA* mutant strain was cultured with supplemental Cu (0, 1, 5, or 10 µM) with or without co-supplemental Mn (0 or 5 µM) for *t* = 8 h. SodA activity was evaluated using an in-gel assay and total protein was evaluated with Coomassie staining. A representative gel from *N =* 3 independent replicates is shown. The activity of purified SodA loaded with either Mn (SpSodA-Mn) or Fe (SpSodA-Fe) was measured in parallel as controls.

### Co-supplementation with Zn or Mn, but not Fe, partially alleviates Cu stress

To further examine the relationship between Cu stress and Zn, Mn, or Fe homeostasis, the Δ*copA* mutant strain was cultured in the presence of Cu and co-supplemental Zn, Mn, or Fe. Figure 5 shows that supplemental Zn or Mn, but not Fe, partially rescued growth of the Δ*copA* mutant strain in the presence of Cu. Since Cu stress in GAS is associated with decreased production of lactate^18^, we also examined whether co-supplemental Zn, Mn, or Fe restores production of this fermentation end-product. Figure 6 shows that co-supplemental Zn or Mn, but not Fe, partially increased total lactate levels secreted by Cu-treated Δ*copA* cells.

**Figure 5.**
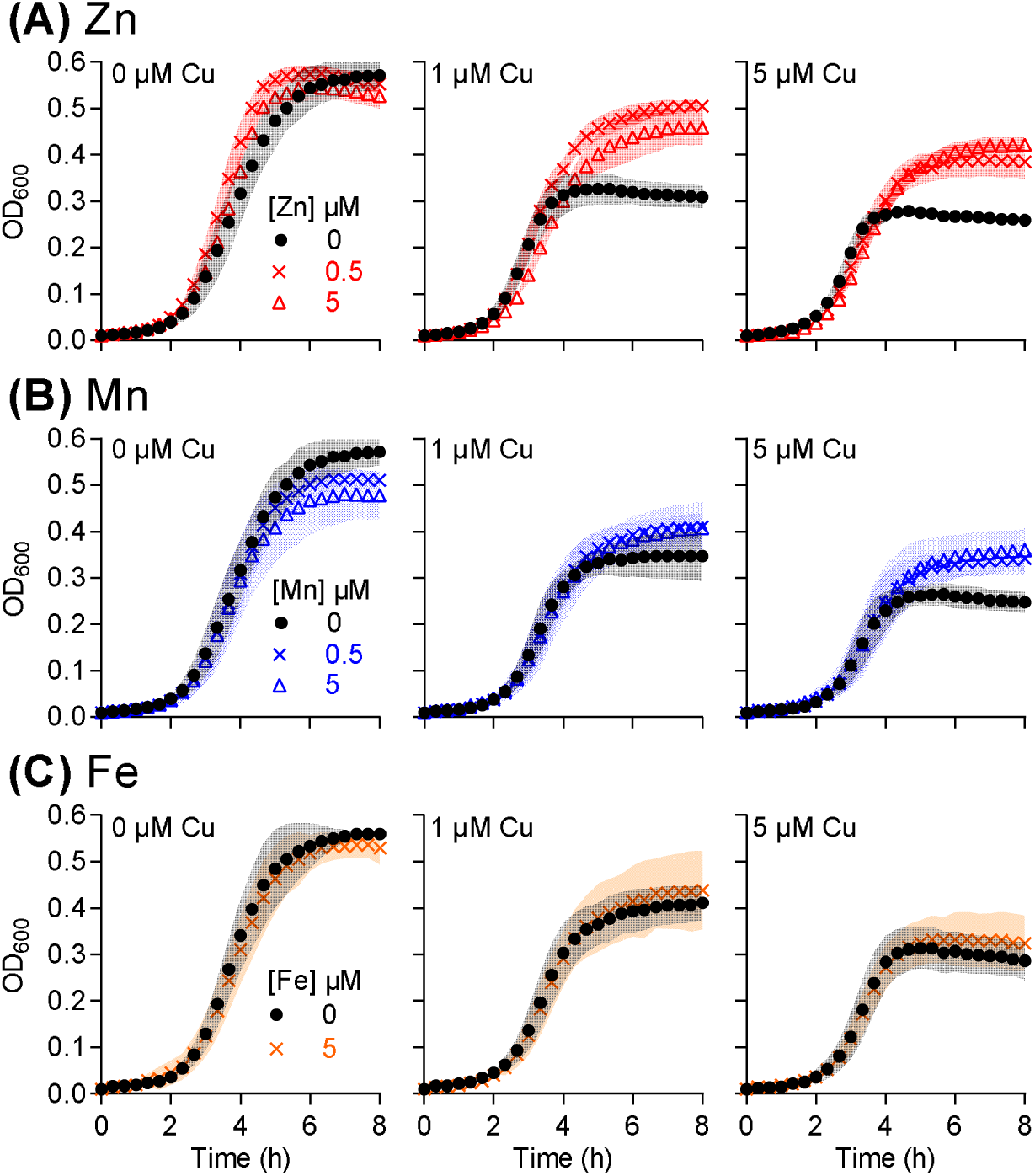
Effects of co-supplemental (A) Zn, (B) Mn, or (C) Fe on bacterial growth. The GAS 5448Δ*copA* mutant strain was cultured with added Cu (0, 1, or 5 µM) with or without added Zn, Mn, or Fe (0, 0.5, or 5 µM) for *t* = 8 h (*N =* 3). Symbols represent means. Shaded regions represent SD. Co-supplemental Zn improved growth (*P* = 0.0006, <0.0001, and <0.0001, respectively for 0, 1, and 5 µM Cu). Mn also improved growth (*P* = 0.44, 0.73, and <0.0001, respectively, for 0, 1, and 5 µM Cu) but to a lesser extent than did Zn. By contrast, Fe had no effect (*P* = 1.0, 0.97, and 0.55, respectively for 0, 1, and 5 µM Cu).

**Figure 6.**
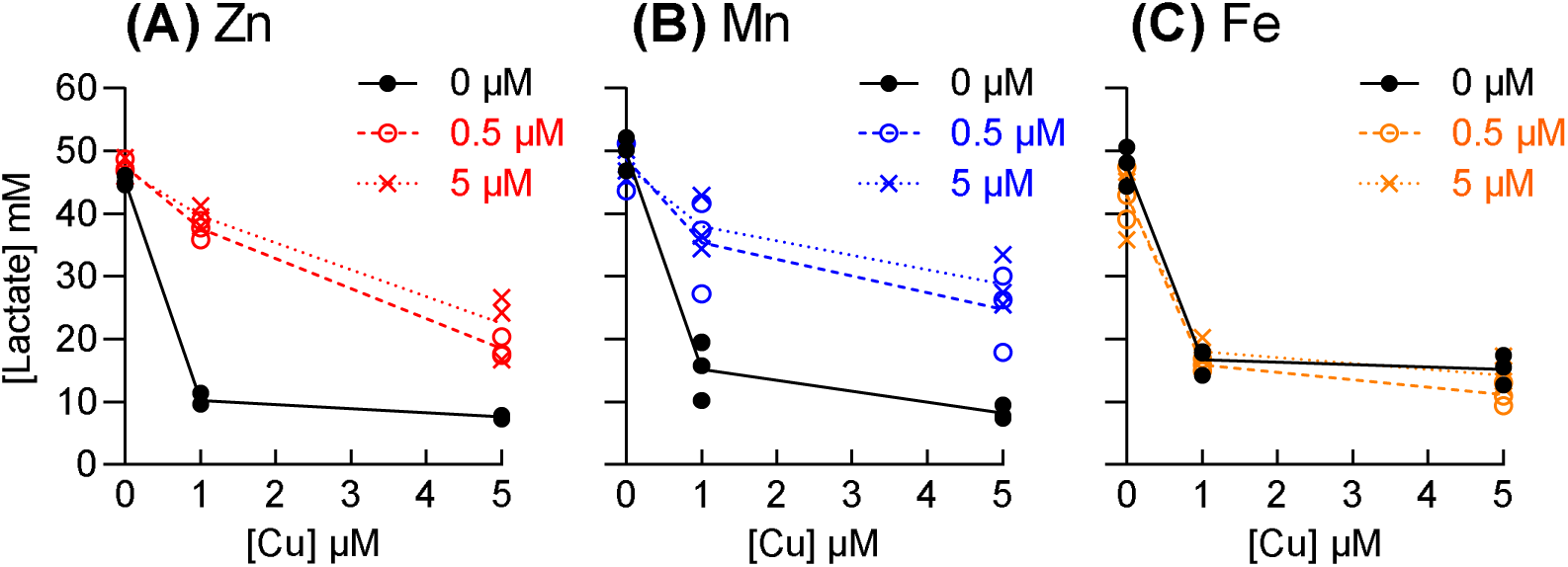
Effects of co-supplemental (A) Zn, (B) Mn, or (C) Fe on lactate levels. The GAS 5448Δ*copA* mutant strain was cultured with added Cu (0, 1, or 5 µM) with or without added Zn, Mn, or Fe (0, 0.5, or 5 µM) for *t* = 24 h (*N* = 3). Amounts of secreted lactate were measured in the spent culture media. Data from individual independent replicates are shown. Lines indicate means. Cu treatment led to a decrease in lactate levels (*P* < 0.0001). This effect was rescued by Zn or Mn (*P* <0.0001 or *P* = 0.0007, respectively) but not Fe (*P =* 0.46).

The absence of a detectable effect by co-supplemental Fe, combined with the lack of a clear relationship between Cu treatment and total Fe metal levels or the expression of *siaA* and *fhuA*, suggests that Cu stress in the Δ*copA* mutant strain is not Fe-dependent, at least under our experimental conditions. For the purposes of this work, the relationship between Cu stress and Fe homeostasis was not investigated further. By contrast, our data clearly hint at a link between Cu stress and Zn or Mn homeostasis, which was explored in more details below.

### Co-supplementation with Zn or Mn does not suppress cellular Cu levels and availability

The simplest model that can explain the protective effects of Zn or Mn during Cu stress is that each metal suppresses cellular Cu levels and/or availability. To test this model, we first measured total cellular Cu levels in Δ*copA* cells that were grown in the presence of Cu and co-supplemental Zn or Mn. The results indicated that neither co-supplemental Zn nor Mn reduced total cellular Cu levels in Δ*copA* cells (Figure 7A). To measure cellular Cu availability, we assessed expression of the Cu-inducible *copZ* gene. We have established previously that deletion of the *copA* gene does not affect Cu-dependent expression of *copZ*^18^, which is immediately downstream from *copA*. As shown in Figure 7B, co-supplemental Zn or Mn did not perturb Cu-dependent de-repression of *copZ* and, therefore, Cu availability.

**Figure 7.**
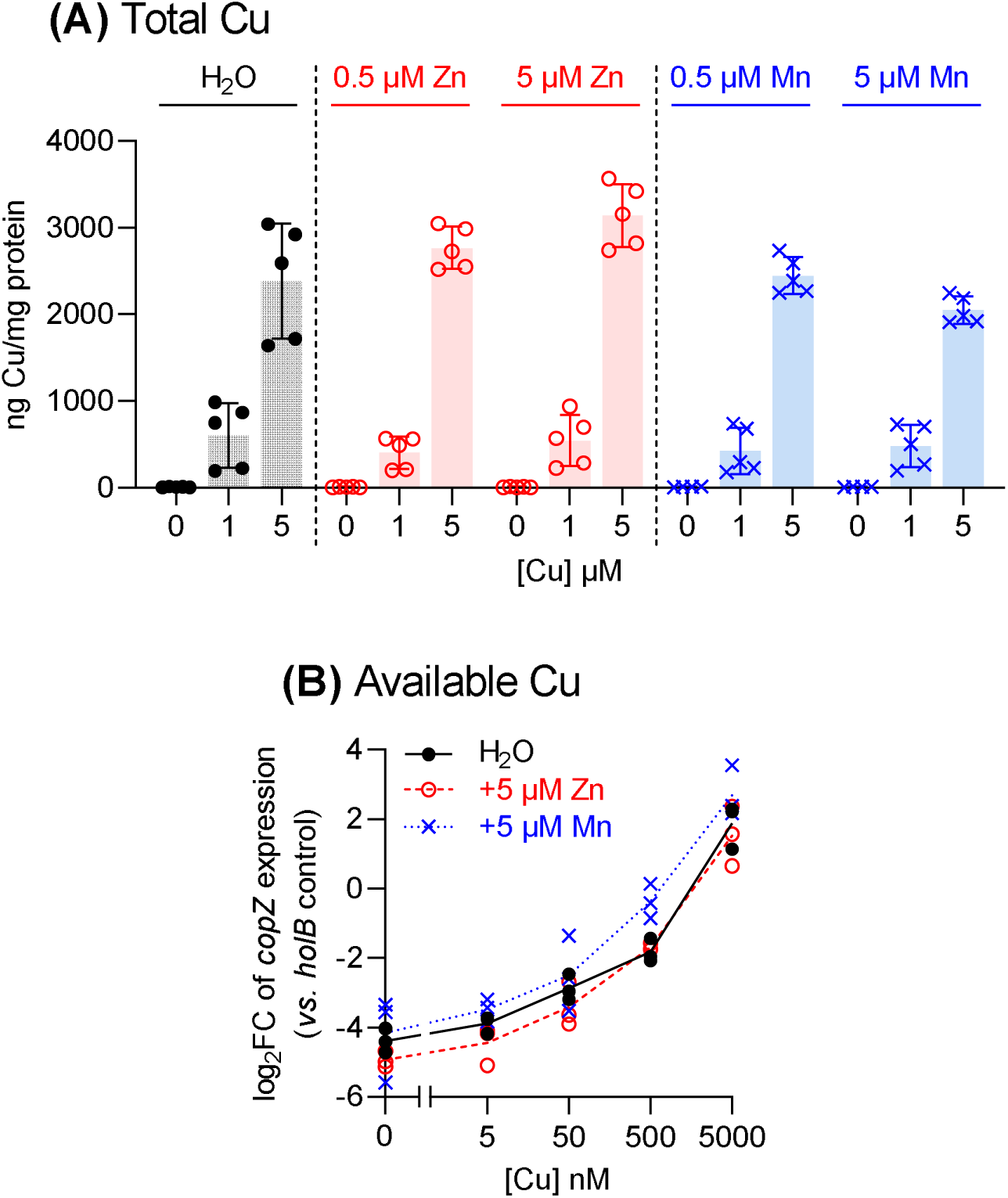
Effects of co-supplemental Zn or Mn on cellular levels of (A) total Cu and (B) available Cu. **(A)** The GAS 5448Δ*copA* mutant strain was cultured with added Cu (0, 1, or 5 µM) with or without Zn or Mn (0, 0.5, or 5 µM) for *t* = 8 h (*N* = 5). Total cellular levels of Cu were measured by ICP MS and normalised to total cellular protein content. Data from individual replicates are shown. Columns indicate means. Error bars represent SD. Cu treatment led to an increase in cellular Cu levels (*P* < 0.0001). Co-supplemental Zn or Mn had no effect on cellular Cu levels (*P =* 0.92, 0.07, 0.99, or 0.35 respectively, for 0.5 µM Zn, 5 µM Zn, 0.5 µM Mn, and 5 µM Mn). Note that Figure 3A shows the same data for Cu levels without co-supplemental Zn or Mn. **(B)** The GAS 5448Δ*copA* mutant strain was cultured with added Cu (0 – 5000 nM) with or without added Zn or Mn (5 µM each) for *t =* 8 h (*N* = 3). Levels of *copZ* mRNA in these samples were determined by qRT-PCR and normalised to expression of *holB* as the control. The normalised expression levels of *copZ* in Cu-treated samples were then compared to those in untreated controls and plotted as log_2_FC values. Individual replicates are shown. Lines represent means. Neither co-supplemental Zn nor Mn affected Cu-dependent *copZ* expression (*P* = 0.70 and 0.53, respectively).

It can be argued that the high affinity of CopY to Cu, as would be expected for a metal-sensing transcriptional regulator, would render it highly sensitive to changes within the low, homeostatic range of cellular Cu availabilities but insensitive to changes within the high, toxic range. Our previous work shows that high cellular Cu availability within the toxic range led to a reduction in the activity of the ATP-generating, GAPDH enzyme, GapA^18^. GapA is likely mismetalated by the excess cytoplasmic Cu ions, which may bind to the side chains of the catalytic Cys and a nearby His^33^. Figure 8 confirms that GapA activity remained low in Δ*copA* cells that were co-supplemented with Zn or Mn. This observation supports our conclusion that co-supplemental Zn or Mn does not influence Cu availability.

**Figure 8.**
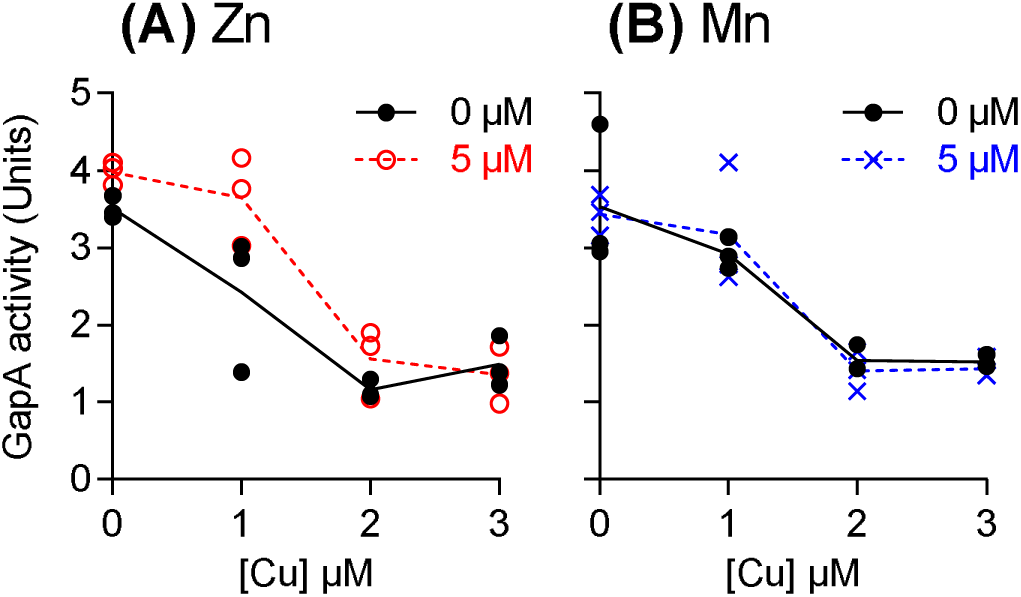
Effects of co-supplemental (A) Zn or (B) Mn on GapA activity. The GAS 5448Δ*copA* mutant strain was cultured with added Cu (0 – 3 µM) with or without added Zn or Mn (5 µM each) for *t* = 8 h (*N =* 3). GapA activities were determined in cell-free extracts. Data from individual replicates are shown. Lines indicate means. Neither Zn nor Mn influenced the effect of Cu on GapA activity (*P* = 0.12 and 0.88, respectively).

### MtsABC or AdcAI/AdcBC does not promote uptake of Cu into GAS

Our results differ from those reported previously in *Staphylococcus aureus*. In *S. aureus*, MntABC, a Mn-importing ABC transporter, is thought to promote uptake of Cu into the cytoplasm^34^. Expression of *mntABC* in *S. aureus* is controlled transcriptionally by the Mn-sensing derepressor MntR^35^. Thus, a decrease in MntABC levels (and activity), either by deletion of the *mntA* gene or by transcriptional repression of the *mntABC* operon in response to Mn co-supplementation, leads to a reduction in cellular Cu levels^34^.

If the MtsABC transporter in GAS and, by extension, the AdcAI/II-AdcBC transporter take up Cu into the GAS cytoplasm, then co-supplemental Mn or Zn may alleviate Cu stress by repressing the transcription of *mtsA*, *adcAI*, or *adcAII* and suppressing Cu uptake *via* their protein products. Similarly, mis-repression of *mtsA*, *adcAI*, or *adcAII* genes by excess Cu potentially suppresses further Cu uptake *via* these transporters and self-limits the toxicity of this metal.

We have already shown that co-supplemental Zn or Mn did not suppress Cu levels or availability in Δ*copA* cells (Figure 7). Under these experimental conditions, co-supplemental Zn (5 µM) was confirmed to increase cellular Zn levels and repress expression of *adcAI* independently of Cu (Supporting Figures 3A and 4A). Interestingly, co-supplemental Mn (5 µM) did not repress the expression of *mtsC* independently of Cu (Supporting Figure 4B), even though cellular Mn levels increased more than tenfold (Supporting Figure 3B). Higher amounts of Mn were not examined because they were inhibitory to the Δ*copA* mutant strain, even in the absence of Cu. Thus, the *mtsC* gene remained active in our experiments. Altogether, these data did not sufficiently address the potential role of the Adc and Mts ABC transporters in promoting the uptake of Cu in GAS.

Thus, we examined deletion mutant strains lacking the relevant ABC transporters (Supporting Table 1). Since there is overlap in the function of AdcAI and AdcAII^36,37^, the Δ*adcAI/II* mutant strain lacking both proteins^37^ was used, along with the Δ*adcBC* mutant strain lacking the AdcBC transmembrane domains^37^. The Δ*mtsABC* mutant strain lacking the entire MtsABC transporter^32^ was also assessed. According to the *S. aureus* model, these different GAS mutant strains would take up less Cu and thus become more resistant to Cu stress when compared with the wild type. Contrary to this hypothesis, none of the mutant strains displayed a Cu-resistant phenotype (Figure 9). In fact, the Δ*mtsABC* mutant strain was reproducibly *less* resistant to the inhibitory effects of Cu than was wild-type strain. Complementation of this mutant *in cis* with a functional copy of the *mtsABC* operon restored the wild-type phenotype (Supporting Figure 5).

**Figure 9.**
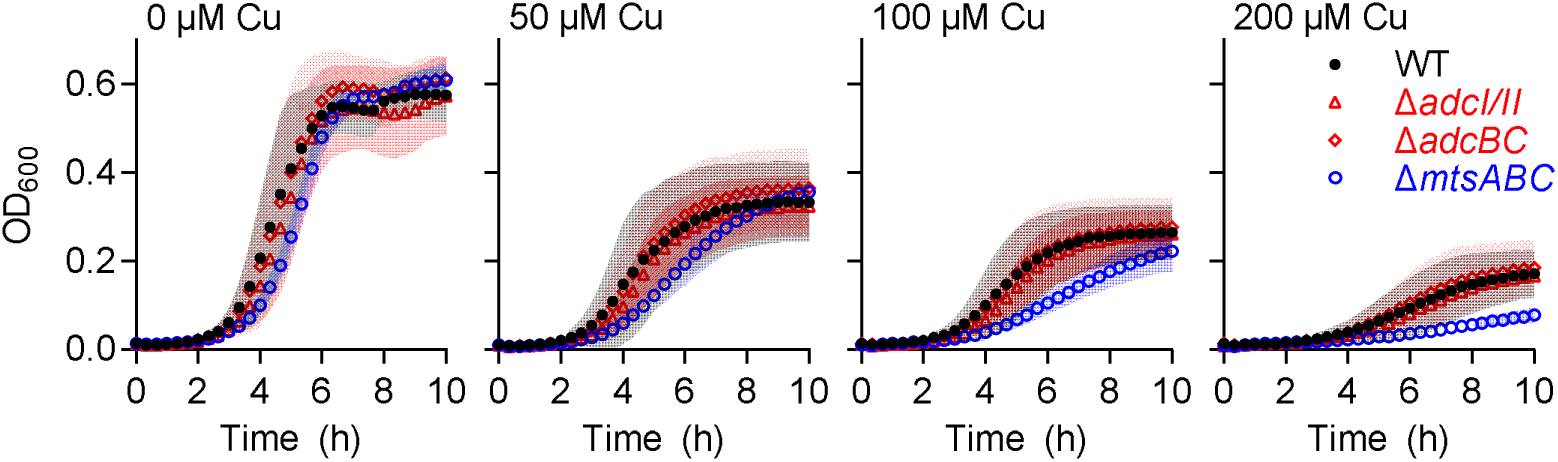
Effects of supplemental Cu on growth of ABC transporter knockout mutant strains. GAS 5448 wild-type and mutant strains were cultured with added Cu (0, 50, 100, or 200 µM) for *t* = 10 h (*N =* 3). Symbols represent the means. Shaded regions represent SD. The different mutations did not affect bacterial growth in the absence of Cu (*P* = 0.88, 0.19, and 0.69, respectively, for Δ*adcAI/II*, Δ*adcBC*, and Δ*mtsABC*). Cu did not affect growth of the Δ*adcAI/II* (*P* = 0.37, 0.52, and 1.0, respectively, for 50, 100, and 200 µM Cu) or Δ*adcBC* (*P* = 0.61, 0.99, and 0.68, respectively, for 50, 100, and 200 µM Cu) mutant strain differently from WT. However, Cu inhibited growth of the Δ*mtsABC* more strongly when compared with the WT (*P* = 0.0009, <0.0001, and <0.0001, respectively, for 50, 100, and 200 µM Cu).

The presence of a functional CopA efflux pump in all the knockout mutant strains used in Figure 9 may mask the inhibitory effects of Cu on bacterial growth. Unfortunately, despite screening thousands of transformants, the double mutant strains Δ*copA*Δ*adcA*/*II*, Δ*copA*Δ*adcBC*, and Δ*copA*Δ*mtsABC* were not obtained.

In the absence of the desired double mutants, we examined whether loss of the transporters in each single mutant strain reduce cellular Cu levels and/or availability. To minimise interference either from Cu efflux by the Cu-inducible CopA pump or from potential aberrant Cu-dependent and - independent changes in the transcription of multiple metal homeostasis genes, the mutant strains were cultured for 8 h in the *absence* of Cu and subsequently exposed to Cu only for 30 min. If an ABC transporter takes up Cu as hypothesised, then we should observe a decrease in cellular Cu levels and/or a time-dependent delay in de-repression of *copA* in the relevant knockout mutant strain when compared with the wild type. As shown in Figure 10A, there was no difference between cellular Cu levels in the wild-type and any mutant strain. Similarly, there was no difference between the expression patterns of the *copA* gene in the different strains (Figure 10B). Therefore, there is currently no experimental evidence to support the uptake of Cu *via* the MtsABC or AdcAI/II-AdcBC transporter in GAS.

**Figure 10.**
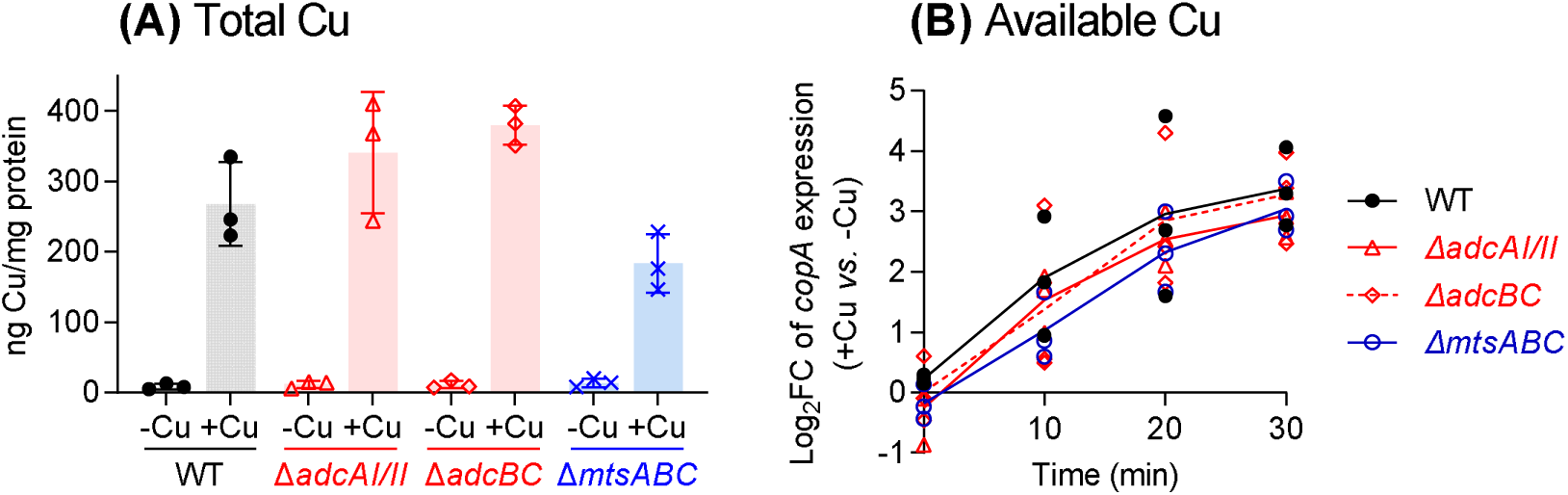
Effects of ABC transporter deletions on cellular levels of (A) total Cu and (B) available Cu. GAS 5448 wild-type and mutant strains were cultured for *t =* 8 h (*N* = 3). **(A)** Each culture was exposed to 5 µM Cu for 30 min. Total cellular Cu levels were measured by ICP MS and normalised to total cellular protein content. Data from individual replicates are shown. Columns indicate means. Error bars represent SD. The different mutations did not influence cellular Cu levels (*P* = 0.36, 0.09, and 0.25, respectively, for Δ*adcAI/II*, Δ*adcBC*, and Δ*mtsABC*). **(B)** Each culture was exposed to 500 nM Cu and sampled at 10 min intervals for 30 min. Levels of *copA* mRNA in each sample was determined by qRT-PCR and normalised to expression of *holB* as the control. The normalised expression levels of *copA* in Cu-treated samples were then compared to those in untreated controls and plotted as log_2_FC values. Individual replicates are shown. Lines represent the means. The absence of the different ABC transporters did not affect the time-dependent expression of *copA* (*P =* 1.0, 0.92, and 0.84, respectively for Δ*adcAI/II*, Δ*adcABC*, and Δ*mtsABC*).

## DISCUSSION

### Zn and Mn homeostasis in GAS are perturbed during Cu stress

This study strengthens our previous observation that growth in the presence of excess Cu leads to mis-regulation of AdcR-and MtsR-dependent metal homeostasis in GAS^18^, although these effects are not associated with detectable changes in cellular Zn or Mn levels. In addition, supplemental Zn or Mn partially alleviates Cu stress, although neither metal appears to influence cellular Cu levels or availability.

To explain these observations, we refer to our previous model^18^, in which the excess cellular Cu binds to the allosteric metal-binding site in AdcR or MtsR, leading to *incorrect* sensing of metals and mis-repression of target genes. The protective effect of co-supplemental Zn or Mn is not inconsistent with this model. Although neither Zn nor Mn interfered with accumulation of cellular Cu (Figure 7), each did increase cellular Zn or Mn levels (Supporting Figure 3) and, at least in the case of Zn, availability (Supporting Figure 4). The additional Zn or Mn may bind directly to their cognate metallosensors, remodel homeostasis of that metal in the cell, and override potential mis-signalling by Cu.

An equally plausible model is that the excess cellular Cu outcompetes Zn or Mn from their binding sites in Zn-and Mn-dependent proteins. Although total cellular levels of Zn or Mn may not change, the unintended dissociation of Zn or Mn from existing binding sites would increase their cellular availability. The latter would again enhance subsequent binding of Zn to AdcR or Mn to MtsR and *correctly* promote transcriptional repression of *adcAI*, *adcAII*, and *mtsC*. In this case, co-supplemental Zn or Mn may promote re-metalation of the mis-metalated Zn or Mn-dependent proteins. Or, co-supplemental Zn or Mn may metalate *other* proteins that allow cells to bypass Cu stress. For example, bacterial growth was restored by co-supplemental Zn or Mn (Figure 5), eventhough the cellular activity of a likely mis-metalation target, namely GapA, remained low (Figure 8). There is currently insufficient biochemical data to distinguish between the different models proposed here.

### Cross-talks between Cu and Zn homeostasis in the bacterial world

Cross-talks between Cu stress and Zn homeostasis have been reported in other bacteria, although the molecular details seem to differ. In *S. pneumoniae*, excess supplemental Zn aggravates (rather than alleviates) Cu stress in a Δ*copA* mutant strain^38^ and in a Δ*czcD* mutant strain lacking the primary Zn efflux transporter^39^. Here, excess Zn in the cytoplasm is thought to bind to the allosteric sensing site of the Cu sensor CopY, stabilise the repressor form of this sensor, and thus suppress transcriptional sensing of Cu^16^. Based on the patterns of *copZ* expression in Figure 9, there is no evidence that Zn perturbs transcriptional Cu sensing in GAS, at least under the experimental conditions employed here, which contain low, non-inhibitory amounts of supplemental Zn.

As another example, Cu treatment in *Salmonella* leads to upregulation (and not downregulation) of Zn uptake genes under the control of the Zn sensor Zur^4^. Whether Cu treatment perturbs Zn levels in this organism has not been reported. In *Acinetobacter baumanii*, supplemental Cu does not perturb Zn levels in wild-type or Δ*copA* mutant strains^40^. However, supplemental Zn does lead to a decrease in cellular Cu levels in the wild-type strain^41^. The molecular mechanism is unclear, but several putative metal transporter genes are differentially regulated in response to Zn, potentially leading to increased efflux or decreased uptake of Cu from the cytoplasm. This scenario resembles that reported in *Escherichia coli*. Supplemental Zn alleviates Cu stress and decreases cellular Cu levels in the *E. coli* wild-type and Δ*cueO* mutant strains^42,43^. In this case, supplemental Zn promotes mis-activation of the *cusCFBA* operon encoding an RND-family Cu efflux transporter, and thus a lowering of cellular Cu^43^. As stated earlier, our work found no evidence that low levels of supplemental Zn perturb transcription of Cu homeostasis genes in GAS.

Similar to our findings, growth of Cu-treated Δ*copA* mutant strains of *S. pneumoniae* is improved by co-supplementation with Mn^38^. The excess Cu in this organism is thought to inhibit the Mn-dependent ribonucleotide reductase NrdF. Therefore, co-supplementation with Mn would presumably restore NrdF activity^38^. Cu may similarly inhibit NrdF in GAS. However, loss of NrdF activity is likely only a minor component of Cu stress in GAS, since co-supplementation with Mn is less protective than co-supplementation with Zn (*cf.* Figure 6). In contrast with our findings, exposure to Cu leads to upregulation of the *mtsABC* operon in a wild-type strain of *Streptococcus agalactiae* and an increase in cellular Mn levels in a Δ*copA* mutant strain^44^. The mechanism behind this observation is yet to be determined.

The apparent differences in the nature and outcome of the abovementioned cross-talks may reflect inherent differences in the biochemistry of the different metal homeostasis systems in the different organisms. Equally, they may reflect differences in experimental design and setup (*e.g.* growth media, growth stage, concentrations of metals, and/or exposure times to metals), leading to different degrees of Cu stress and/or protection by other metals. For instance, our present study detected a link between Cu and *mtsC* only when the Δ*copA* mutant strain of *S. pyogenes* was cultured beyond 4 h of growth. This time-dependent mis-repression of gene expression is likely associated with the time-dependent depletion of intracellular glutathione and, therefore, time-dependent increase in intracellular Cu availability.

### Do ABC transporters promote Cu uptake into GAS?

Our work further suggests that neither the Zn-importing AdcAI/II-AdcBC transporter nor the Mn-importing MtsABC transporter promotes uptake of Cu into GAS. To take up a metal ion, the extracytoplasmic solute binding protein (SBP) domain captures its cognate metal ion and subsequently releases this metal to the metal-binding site in the permease domain. In turn, the permease internalises the metal ion into the cytoplasm and this action is powered by ATP hydrolysis by the nucleotide-binding domain. Unpublished studies in our laboratory suggest that AdcAI, the Zn-binding SBP from GAS, binds Cu(II) more tightly than it binds Zn(II). Although the AdcAII SBP from GAS has not been biochemically characterised, the homologue from *S. pneumoniae* has also been reported to bind Cu(II)^45^. Likewise, the Mn-binding MtsA SBP from GAS binds Cu(II)^46^, as does PsaA, the MtsA homologue from *S. pneumoniae*.

Our data suggest that the bound Cu(II) in any of the above SBPs is not transferred to the metal-coordinating site in the partner permease and subsequently internalised into the cytoplasm. There is evidence that an SBP does not load the permease with non-cognate metal ions, a result of incompatible coordination chemistry between the partners. For example, extracellular Zn competitively inhibits Mn uptake *via* PsaABC in *S. pneumoniae*^8,9,47^. The permeases PsaB (which imports Mn) and AdcB (which imports Zn) in this organism possess the same, conserved metal coordination site^48^, suggesting that PsaB should be competent to receive Zn from PsaA. However, while PsaC efficiently releases the bound Mn to PsaB, it does not release bound Zn^47^. Whether Zn can access the metal-binding site in PsaB directly, without the PsaA SBP, is unknown. Similarly, whether excess extracellular Cu can bind directly to the metal-binding site in the Zn-importing permease AdcB or the Mn-importing permease MtsC and become subsequently internalised into the cytoplasm is unknown. Our data do not support this hypothesis, but direct biochemical evidence, for instance *via* metal transport assays of purified transporters, remains to be obtained.

## AUTHOR CONTRIBUTIONS

KD initiated and designed the research. KD also had overall responsibility for conceptualizing and coordinating the programme. KD and LS measured bacterial growth, GapA activity, and cellular metal levels. JB and KD measured gene expression levels. EM and KW measured SodA activity. SF and YH carried out literature review and preliminary experiments leading to the final work shown here. KD drafted the manuscript with input from SF and YH. KD, YH, and EM prepared the figures. KD, SF, YH, EM, and KW edited the manuscript. All authors approved the final version. JB, SF, JH, EM, LS have contributed equally to this work. The order of names in the author list was decided by an on-line random list generator.

## ACKNOWLEDGEMENT

We thank Dr Peter Chivers (Department of Biosciences, Durham University) for helpful discussions related to this project. Dr Christopher Ottley (Department of Earth Sciences, Durham University) and Dr Emma Tarrant (Department of Biosciences, Durham University) provided technical assistance with ICP MS. The Δ*mtsABC*, Δ*adcAI/II*, and Δ*adcBC* mutant strains are kind gifts from Dr Andrew Turner, Dr Cheryl-lynn Ong, Prof. Alastair McEwan, and Prof. Mark Walker (School of Chemistry and Molecular Biosciences, The University of Queensland). We also acknowledge the anonymous reviewers of the earlier version of this work for helping us improve this study by prompting us to take a second look at our ICP MS data.

