## Supporting Information for "Mis-regulation of Zn and Mn homeostasis is a key phenotype of Cu stress in *Streptococcus pyogenes*"

**Supporting Table 1.** List of GAS strains used in this study.

| Bacterial strains | Description | Reference |
| --- | --- | --- |
| 5448 (WT) | Invasive M1T1 strain | (1) |
| 5448Δ*copA* | Δ*copA*::*kan* deletion mutant | (2) |
| 5448Δ*adcAI/II* | Δ*adcA*::*kan* Δ*adcAII*::*aad* double deletion mutant | (3) |
| 5448Δ*adcBC* | Δ*adcBC*::*kan* deletion mutant | (3) |
| 5448Δ*mtsABC* | Δ*mtsABC::aphA-3* deletion mutant | (4) |
| 5448Δ*mtsABC::mtsABC+* | *ΔmtsABC* marker rescue mutant | (4) |

**Supporting Table 2.** List of primers used in this study. Primers were designed using the genome sequence of *S. pyogenes* MGAS5005 as template (NCBI GenBank Accession No CP000017.2).

| **Amplicon name** | **Primer name** | **Sequence** | **Locus Tag of target gene** | **Amplicon size (bp)** |
| --- | --- | --- | --- | --- |
| adcAI | adcAI-qPCR-F | GGAACTGGTAACATGCTCTTGG | M5005_Spy0543 | 109 |
|  | adcAI-qPCR-R | CATGGTTGTGTCCTTCTTCGC |  |  |
| adcAII | adcAII-qPCR-F | AGGTTGATGTGTTTGAAGCG | M5005_Spy1711 | 106 |
|  | adcAII-qPCR-R | GGGTCATAAAGTGTCGCAGG |  |  |
| adcC | adcC-qPCR-F | CCATCCACCGTTTACGAGTTTG | M5005_Spy0078 | 99 |
|  | adcC-qPCR-R | GCTTGCTTGCACATGCTCTTC |  |  |
| copA | copA-qPCR-F | TGGTCTCAGGTCTTGTGGTC | M5005_Spy1405 | 97 |
|  | copA-qPCR-R | GTTGCAGAAATGGAGGAAGCC |  |  |
| copZ | copZ-qPCR-F | GCAATCGGTCCAGGTAAATTTGG | M5005_Spy1404 | 92 |
|  | copZ-qPCR-R | TGGTATCCTTCAAAGCACGC |  |  |
| holB | holB-qPCR-F | CATAGTAATCGTAGTGGGTTTCGC | M5005_Spy1835 | 91 |
|  | holB-qPCR-R | CATTTGAAGTCTGCTTAGAACGTG |  |  |
| mtsC | mtsC-qPCR-F | ACTCTGTCATTAAAGGAGATACGGC | M5005_Spy0370 | 100 |
|  | mtsC-qPCR-R | AAGTCCGTCGAACTATTGGC |  |  |
| siaA | siaA-qPCR-F | TGTTGAGGGCATGTACCAGTC | M5005_Spy1528 | 93 |
|  | siaA-qPCR-R | TAGTCCTGGTATTGCTGGCG |  |  |
| fhuA | fhuA-qPCR-F | TTCTTACGGACGTTTTCCCC | M5005_Spy0324 | 104 |
|  | fhuA-qPCR-R | CATAGGCCATGACATTGGTGG |  |  |
| gapA | gapA-check-F | GTAGTTAAAGTTGGTATTAACGG | M5005_Spy0233 | 1008 |
|  | gapA-check-R | TTTAGCAATTTTTGCGAAGTACTCA |  |  |

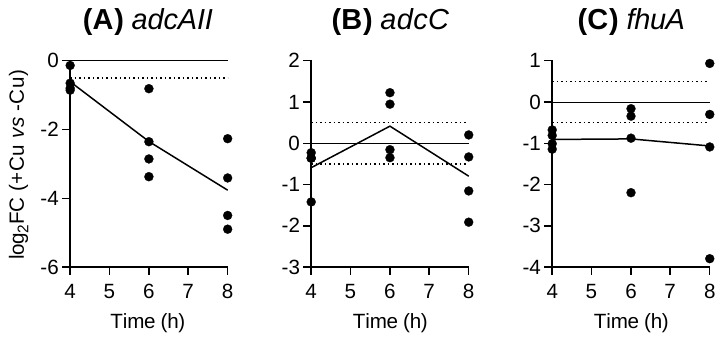

**Supporting Figure 1. Effects of Cu treatment on expression levels of (A) *adcAII*, (B) *adcC*, and (C) *fhuA*.** The GAS 5448∆*copA* mutant strain was cultured with or without added Cu (5 µM) for *t =* 4, 6, or 8 h (*N* = 4). mRNA levels of target genes in Cu-supplemented cultures (+Cu) were determined by qRT-PCR and normalised to those in the corresponding unsupplemented samples (‑Cu) that were cultured for the same time periods. Dotted horizontal lines represent the sensitivity limit of the assay (log_2_FC = ± 0.5). Data from individual replicates are shown. Lines indicate means. Expression levels of *adcAII* (*P* = 0.012) were time-dependent but not those of *adcC* (*P* = 0.094) or *fhuA* (*P* = 0.98).

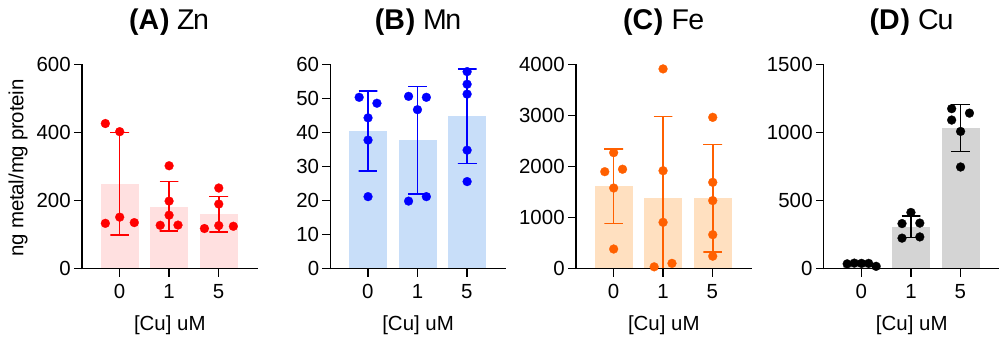

**Supporting Figure 2. Effects of Cu treatment on cellular levels of (A) Zn, (B) Mn, (C) Fe, and (D) Cu, sampled at *t* = 4 h.** The GAS 5448∆*copA* mutant strain was cultured with supplemental Cu (0, 1, or 5 µM) for *t* = 4 h (*N* = 5). Total cellular levels of all metals were measured by ICP MS and normalised to total cellular protein content. Cu treatment influenced cellular levels of Cu (*P* < 0.0001) but not those of Zn, Mn, or Fe (*P* = 0.12, 0.11, or 0.86 respectively).

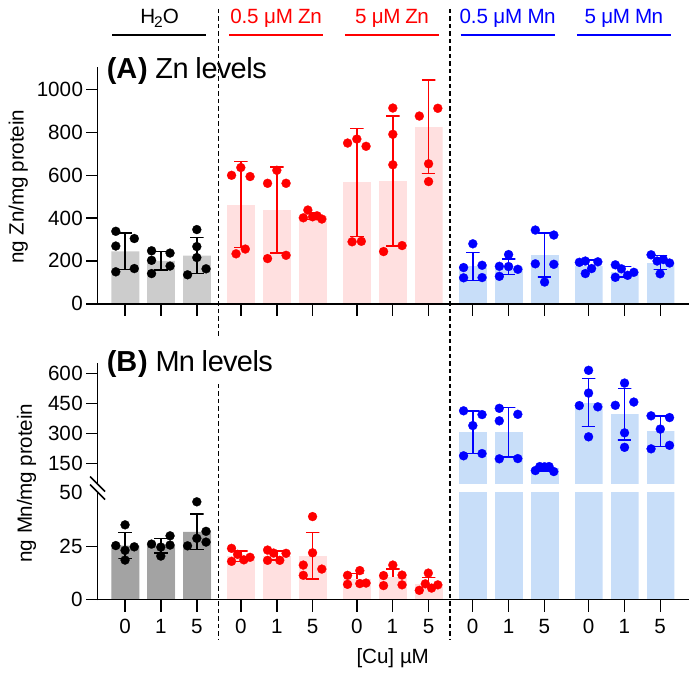

**Supporting Figure 3. Effects of co-supplemental Zn and Mn on cellular levels of (A) Zn and (B) Mn.** The GAS 5448∆*copA* mutant strain was cultured with added Cu (0, 1, or 5 µM) with or without Zn or Mn (0, 0.5, or 5 µM) for *t* = 8 h (*N* = 5). Total cellular levels of Zn and Mn were measured by ICP MS and normalised to total cellular protein content. Note that the H_2_O data are also shown in Figure 3. Data from individual replicates are shown. Columns indicate means. Error bars represent SD. Co-supplemental Zn led to an increase in cellular Zn levels (*P* = 0.0007 and <0.0001, respectively, for 0.5 and 5 µM Zn) but not Mn levels (*P* = 1.0 and 0.86, respectively, for 0.5 and 5 µM Zn). Co-supplemental Mn led to an increase in cellular Mn levels (*P* <0.0001 each for 0.5 and 5 µM Mn) but not Zn levels (*P* = 0.93 and 0.75, respectively, for 0.5 and 5 µM Mn).

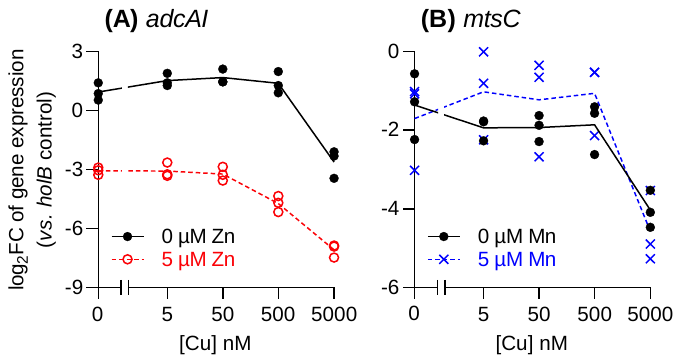

**Supporting Figure 4. Effects of co-supplemental Zn or Mn on Cu-dependent expression of (A) *adcAI* or (B) *mtsC*.** The GAS 5448∆*copA* mutant strain was cultured with added Cu (0 – 5000 nM), with or without added Zn or Mn (5 µM each) for *t* = 8 h (*N* = 3). Levels of *adcAI* or *mtsC* mRNA in these samples were determined by qRT-PCR and normalised to expression of *holB* as the control. Growth in the presence of Cu alone (*P* < 0.0001) or Zn alone (*P* < 0.0001) suppressed the expression of *adcAI*. Growth in the presence of Cu alone suppressed *mtsC* expression (*P* < 0.0001). However, growth in the presence of Mn alone did not suppresss *mtsC* expression (*P* = 0.34).

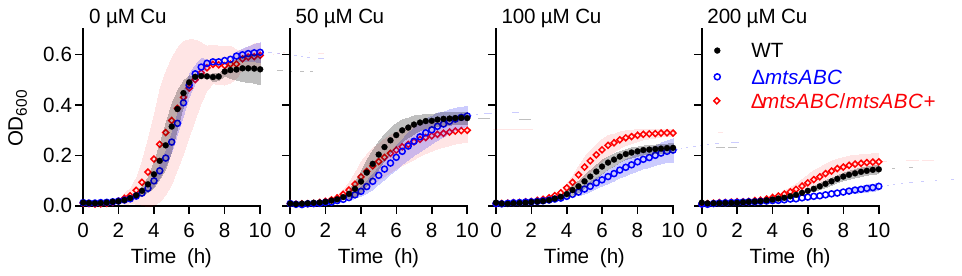

**Supporting Figure 5. Effects of Cu on bacterial growth of the ∆*mtsABC* mutant strain.** The GAS 5448 wild-type (filled circles), ∆*mtsABC* mutant (empty circles), and ∆*mtsABC*/*mtsABC^+^* complemented mutant (empty diamonds) strains were cultured with added Cu (0, 100, 200, 300, or 400 µM) for *t* = 10 h (*N =* 3).

(2)	Stewart, L. J.; Ong, C. Y.; Zhang, M. M.; Brouwer, S.; McIntyre, L.; Davies, M. R.; Walker, M. J.; McEwan, A. G.; Waldron, K. J.; Djoko, K. Y. Role of Glutathione in Buffering Excess Intracellular Copper in Streptococcus Pyogenes. *mBio* *11* (6), e02804-20. https://doi.org/10.1128/mBio.02804-20.

(3)	Ong, C. Y.; Berking, O.; Walker, M. J.; McEwan, A. G. New Insights into the Role of Zinc Acquisition and Zinc Tolerance in Group A Streptococcal Infection. *Infection and Immunity* **2018**, *86* (6). https://doi.org/10.1128/IAI.00048-18.

(4)	Turner, A. G.; Djoko, K. Y.; Ong, C. Y.; Barnett, T. C.; Walker, M. J.; McEwan, A. G. Group A Streptococcus Co-Ordinates Manganese Import and Iron Efflux in Response to Hydrogen Peroxide Stress. *Biochemical Journal* **2019**, *476* (3), 595–611. https://doi.org/10.1042/BCJ20180902.
